# Cholinergic heterogeneity facilitates synchronization and information flow in a whole-brain model

**DOI:** 10.1101/2025.10.28.685048

**Authors:** Leonardo Dalla Porta, Jan Fousek, Alain Destexhe, Maria V. Sanchez-Vives

## Abstract

The human brain displays substantial regional variability in molecular, anatomical, and physiological organization. Yet, how this heterogeneity shapes large-scale neuronal dynamics remains poorly understood. To address this question, we employed a biologically informed whole-brain computational model capable of generating distinct brain states, from awake-like to sleep-like regimes. Our model was constrained by empirical human structural connectivity and spatial maps of cholinergic receptor gene expression, thereby embedding regional neuromodulatory variability into a macroscopic framework. We found that incorporating cholinergic heterogeneity had a significant impact on brain dynamics: it not only facilitated network synchronization but also enhanced information flow between brain regions. Furthermore, we addressed a particularly intricate dynamic regime characterized by the coexistence of localized sleep-like activity within otherwise awake-like states. We showed that the emergence of these slow waves was a byproduct of both regional levels of neuronal adaptation and structural connectivity. In summary, our findings highlight the critical role of molecular and anatomical heterogeneity in shaping global brain dynamics, suggesting new avenues for linking microscale diversity to macroscale function.

## Introduction

From wakefulness to sleep, the brain exhibits a rich repertoire of spatiotemporal dynamics that underlie behavior and cognition ^1–3^. These dynamics arise from the interplay of factors across multiple scales, including structural connectivity, cellular physiology, and neuromodulatory signaling, each influencing both normal cortical function and susceptibility to pathology ^4^. Yet, the extent to which regional heterogeneity modulates large-scale brain dynamics remains an open question in contemporary neuroscience, particularly across distinct brain states. Neuromodulators and neurotransmitters are well known to shape the functional and dynamical properties of cortical circuits ^5^, but their distribution across the cortex is far from uniform. Instead, these molecular signals form spatially distinct hubs along the cortical hierarchy ^4,6,7^. Despite this, how such molecular and regional heterogeneity gives rise to global brain dynamics and how this relationship depends on brain state remains poorly understood.

Computational models of the human brain have attempted to address this question by simulating large-scale neural dynamics ^8,9^. However, most of these models have focused on the structural properties, largely due to the earlier lack of detailed, molecular-level data. Recent advances in neuroimaging and transcriptomic techniques have advanced the field by enabling the construction of more biologically realistic heterogeneous brain models ^6,7^. Computational models incorporating these region-specific features, such as the T1w:T2w ratio or estimates of excitation-inhibition balance, have demonstrated improved performance in capturing empirical functional dynamics ^10–12^.

Despite recent progress, the influence of molecular heterogeneity on global brain dynamics remains underexplored. Acetylcholine (ACh), for example, is known to play a key role in modulating brain states by influencing spike-frequency adaptation and cortical excitability ^5,13,14^. Cortical ACh levels vary with brain state and exhibit spatial heterogeneityreflecting non-uniform cholinergic projections ^15,16^. Notably, reduced cholinergic input correlates with spatially localized slow wave activity during REM sleep^17^, raising broader questions about the role of cholinergic heterogeneity in generating sleep-like activity under both physiological and pathological conditions, such as attentional lapses or following brain lesions ^18–20^.

To address these questions, we developed a large-scale, biophysically grounded cortical circuit model that incorporates regional heterogeneity informed by the spatial distribution of cholinergic receptors. By simulating different brain states, we found that regional heterogeneity significantly influences large-scale brain dynamics when compared to its homogenous counterpart. Beyond facilitating network coordination through synchronization, heterogeneity was shown to enhance inter-regional communication, supporting more complex and flexible patterns of activity. Furthermore, we demonstrated that spatially localized slow waves emerge as a byproduct of both regional variations in neuronal adaptation and the underlying structural connectivity. In summary, our findings underscore the critical role of both molecular and anatomical heterogeneity in shaping global brain dynamics and provide a novel framework for linking microscale diversity to macroscale brain function.

## Results

Inspired by empirical evidence demonstrating the modulatory effects of acetylcholine on local neuronal circuits and ionic channels ^13^, we investigated here how spatial heterogeneity in cholinergic receptor density influences large-scale brain activity. Using a biophysically grounded whole-brain network model, our goal was to understand how regional variability in cholinergic modulation shapes the spatiotemporal dynamics of the cortex ^3^. In particular, we examined how this heterogeneity impacts asynchronous and synchronous activity patterns, dynamics that closely resemble awake-like and sleep-like brain states, respectively.

### Framework overview

Our whole-brain modeling framework integrates three key components: node dynamics, network connectivity, and regional heterogeneity (Fig. 1). Each brain region (node) was modeled using the Adaptive Exponential Mean-Field (AdEx-MF) model, which captures the interaction between excitatory (*v*_*E*_) and inhibitory (*v*_*I*_) neuronal populations, along with an adaptation variable (*W*) ^21^. W represents activity-dependent processes that modulate neuronal excitability, such as activity-dependent K^+^ currents ^13,22,23^. The dynamic balance between excitation, inhibition, and adaptation allows each node to exhibit either asynchronous (awake-like) or synchronous (sleep-like) dynamics, depending on the level of adaptation. Specifically, increasing neuronal adaptation drives the system toward a synchronous pattern of activity, while lower adaptation promotes irregular, desynchronized activity (Fig. 1, top panel) ^21,24^.

**Fig. 1.**
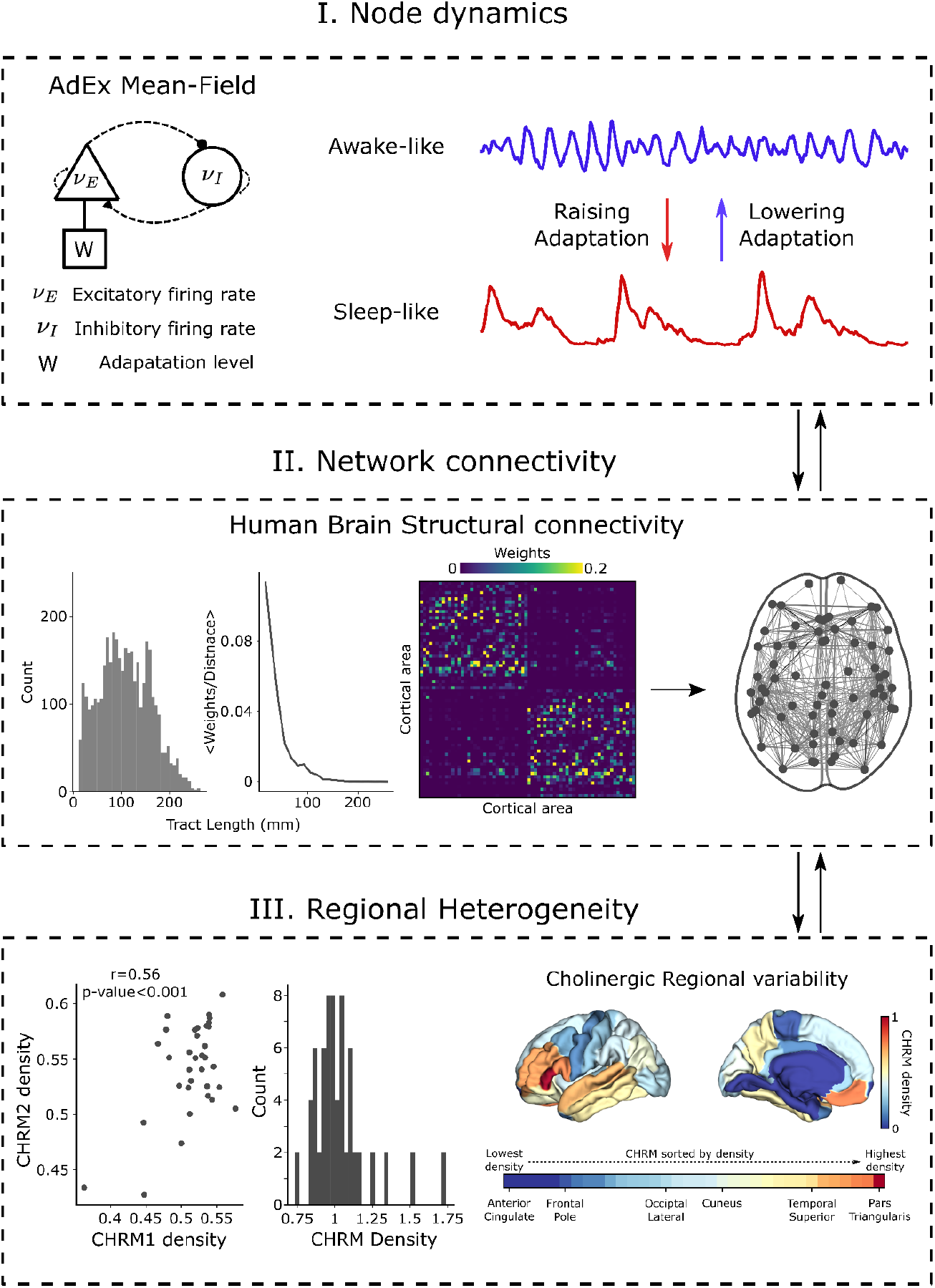
Overview of the whole-brain network model framework. Top panel: Each brain region (node) was modeled using the Adaptive Exponential Mean-Field (AdEx-MF) model. In this model, excitatory (E) and inhibitory (I) neuronal populations are coupled together along with an adaptation variable (W). By varying the adaptation strength, the model can exhibit distinct dynamical regimes: low adaptation leads to asynchronous, awake-like activity, whereas high adaptation produces synchronous, sleep-like activity. Middle panel: Network connectivity was defined using empirical structural connectivity data derived from human diffusion MRI tractography. The brain was parcellated in 68 cortical regions according to the Desikan-Killiany atlas, and connection strength and transmission delays were estimated from diffusion tensor imaging (DTI) data. Inter-regional connectivity was set as excitatory. The preprocessed data was obtained from ^25^. Bottom panel: Regional heterogeneity was introduced in the form of muscarinic acetylcholine receptor density. Gene expression levels of M1 (CHRM1) and M2 (CHRM2) receptor subtypes were highly correlated, and their average (CHRM density) was used to modulate the local adaptation parameter (see Methods for details). Gene expression profiles were obtained from the Allen Human Brain Atlas (AHBA) and processed using the ABAGEN toolbox ^6,31^.

To define network connectivity, we used empirical structural connectivity (SC) data derived from human diffusion MRI tractography (Fig. 1, middle panel; see Methods) ^25,26^. The brain was parcellated according to the Desikan-Killiany atlas (68 cortical regions), and inter-regional connections were modeled as excitatory. Connection strengths (weights) and transmission delays were based on estimates of fiber density and tract length, respectively, obtained from diffusion tensor imaging (DTI) data ^27^. This preprocessed dataset was obtained from ^25^.

Finally, regional heterogeneity was implemented by incorporating spatial variation in cortical receptor density (Fig. 1, bottom panel). Specifically, we used the average expression levels of muscarinic acetylcholine receptor subtypes M1 (CHRM1) and M2 (CHRM2), the two most abundant muscarinic receptors in the human cortex ^6,28,29^. Gene expression profiles were obtained from the Allen Human Brain Atlas (AHBA) and processed using the ABAGEN toolbox ^6,30,31^. These regional expression values were used to modulate the adaptation parameter in each node, thus linking neuromodulatory heterogeneity to local excitability. Altogether, this approach allowed us to construct a biophysically grounded, spatially heterogeneous virtual brain model (Fig. 1).

### From asynchronous to synchronous dynamics in a large-scale network model

Following our framework previously described, we next investigated the whole-brain network dynamics as a function of the adaptation level (b; Fig. 2). At low adaptation levels (<b>=12 pA, <> indicates average over all brain areas), the system exhibited weak inter-regional correlations, as illustrated by the functional connectivity (FC) matrix, resembling the desynchronized activity typically associated with awake-like states (Fig. 2A-D). Accordingly, temporal fluctuations in this regime were characterized by low-amplitude, high-frequency activity, which gave rise to irregular oscillations in the range of ∼10Hz, also accompanied by a suppression of the low-frequency content (Fig. 2B-C). Consequently, this desynchronized state was dominated by irregular fluctuations, resulting in low correlation values across brain regions (Fig. 2D).

**Fig. 2.**
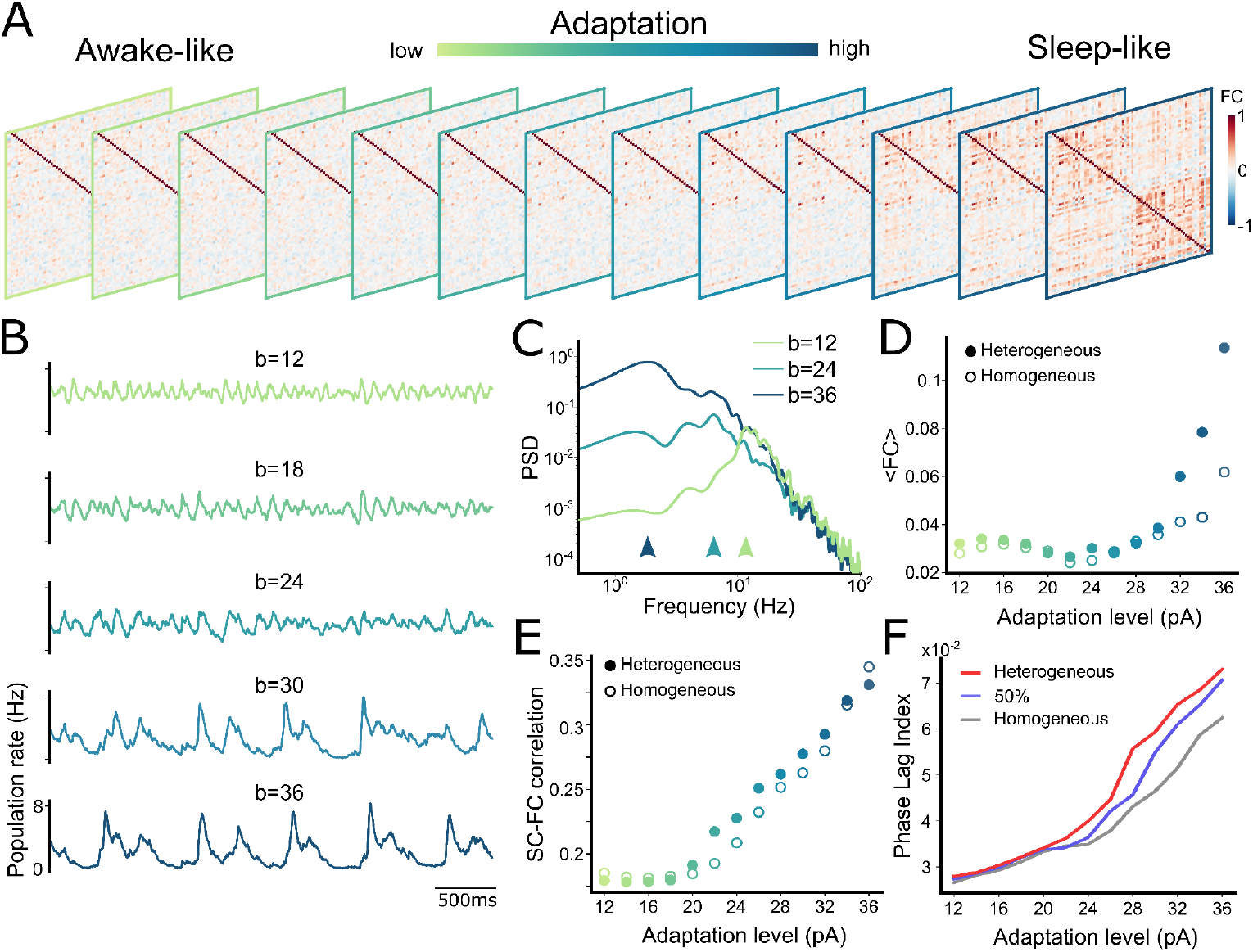
Heterogeneous whole-brain dynamics from asynchronous to synchronous dynamics. (A) Functional connectivity (FC; Pearson correlation) matrices as a function of adaptation levels. (B) Representative time series for different adaptation values (b in pA units). (C) Power spectrum density (PSD) for three representative adaptation levels illustrating awake-like (<b>=12 pA), intermediate (<b>=24 pA), and sleep-like (<b>=36 pA) dynamics. Arrows indicate the peak frequency at ∼11 Hz, ∼6 Hz, and ∼2 Hz, respectively. (D) Mean FC as a function of adaptation level for heterogeneous (filled circles) and homogeneous (empty circles) networks. (E) Same as (D) for structure-function (SC-FC) coupling (Pearson correlation between structural and functional connectivity). (F) Phase lag index as a function of adaptation level. Red and gray lines represent heterogeneous and homogenous networks, respectively. Light-purple indicates a partially heterogeneous network (50% heterogeneity; see Methods for details).

As adaptation increased (<b>=24 pA), inter-regional correlations also increased, and locally connected clusters began to emerge, giving rise to structured spatiotemporal patterns (Fig. 2A). In this intermediate regime, network firing rate decreased (from ∼6Hz to ∼4Hz), and the dominant oscillatory frequency slowed down (from ∼10Hz to ∼6Hz), alongside a relative increase in the low-frequency content (Fig. 2B-C). Despite the emergence of localized synchrony, as indicated by the FC matrices, the global network remained weakly correlated (Fig. 2D). We further discuss this intermediate regime in the following section (see *Emergence of localized sleep-like slow waves*).

For higher adaptation levels (<b>=36), the network dynamics were dominated by strongly correlated fluctuations across widespread cortical regions, a dynamic regime reminiscent of SWS and deep anesthesia (Fig. 2A) ^32–34^. In this state, the system exhibited quasi-periodic (∼1Hz) alternations between periods of high-amplitude (∼8Hz) sustained activity (Up states) and periods of network silence (Down states or off-periods), which propagated through the network as travelling waves (Supp. Fig. 1). As a result, low-frequency components dominated the system, leading to a globally synchronized network state (Fig. 2D).

Another important feature of awake- and sleep-like states is their increased structure-function coupling, i.e., the relationship between the underlying anatomical structure and the emergent functional patterns ^35–38^. Our model reproduced this relationship accurately (Fig. 2E). Specifically, for awake-like states (desynchronized activity), the FC was only loosely constrained by the SC. Conversely, during sleep-like states (synchronized activity), the FC patterns were more constrained by the SC. This strengthened FC–SC coupling is thought to reflect a reduced repertoire of dynamic brain states, which is typically higher during wakefulness and significantly lower during sleep and unconscious states ^24,35,36^.

### Regional heterogeneity facilitates large-scale synchronization

In the previous section, we demonstrated that our whole-brain network model, incorporating regional heterogeneity constrained by the spatial distribution of cholinergic receptor density (see Methods), can reproduce key features of both awake-like and sleep-like brain dynamics (Fig. 2). However, a crucial question remains: how does the inclusion of regional heterogeneity impact global brain dynamics? To address this, we compared our heterogeneous model to its homogeneous counterpart, i.e., keeping the adaptation level constant across brain regions, i.e., heterogeneity = 0% (<b> = b).

By comparing FC profiles across adaptation levels, we found that regional heterogeneity facilitated network synchronization; however, only at higher adaptation values (Fig. 2D; compare filled vs. empty circles). For adaptation current levels above 30 pA, the mean FC (measured via Pearson correlation) was significantly higher in the heterogeneous model, indicating that regional variability in adaptation enhances inter-regional coupling under synchronized states. Interestingly, although the heterogeneous network exhibited stronger overall synchronization, the structure–function coupling remained similar (Fig. 2E). That is, the relationship between FC and SC did not significantly differ between the heterogeneous and homogeneous models, suggesting that, despite higher synchronization, the dynamics remained constrained by the underlying anatomical structure.

In our model, synchronous states are characterized by Up and Down dynamics, which propagate through the network as travelling waves (Supp. Fig. 1). To further investigate the impact of regional heterogeneity on network synchronization, we computed the Phase Lag Index (PLI; see Methods). PLI is a metric designed to quantify phase synchronization while minimizing the influence of zero-lag correlations ^39^. Thus, PLI is particularly well-suited for detecting non-trivial phase relationships, such as those expected during traveling wave dynamics.

PLI analysis confirmed that regional heterogeneity indeed enhances network synchronization (Fig. 2F). For adaptation levels above <b>=20 pA, heterogeneous networks exhibited higher synchronization compared to their homogeneous counterparts. This difference became more pronounced at higher adaptation levels, which are associated with stronger global synchronization. Notably, the effect of heterogeneity on synchronization appeared to be gradual: networks with 50% heterogeneity displayed intermediate PLI values between the fully heterogeneous and fully homogeneous cases. Altogether, these results suggest that regional heterogeneity facilitates large-scale network coordination by enhancing phase synchronization across brain regions.

### Information flow is enhanced in large-scale networks with regional heterogeneity

A fundamental aspect of neuronal networks is their ability to receive and transmit information across the connectome, a process supported by both local and long-range connections ^40–42^. While previous studies have shown that information flow is enhanced in local spiking networks with heterogeneous neuronal properties, this effect has been less explored at the large scale ^43–45^. We next assessed the information flow across the connectome with regional heterogeneity and in its homogeneous counterpart. To this end, we briefly stimulated a single cortical area under both awake-like and sleep-like dynamic regimes and computed the transfer entropy (TE) between the stimulated region and all other areas (Fig. 3).

**Fig. 3.**
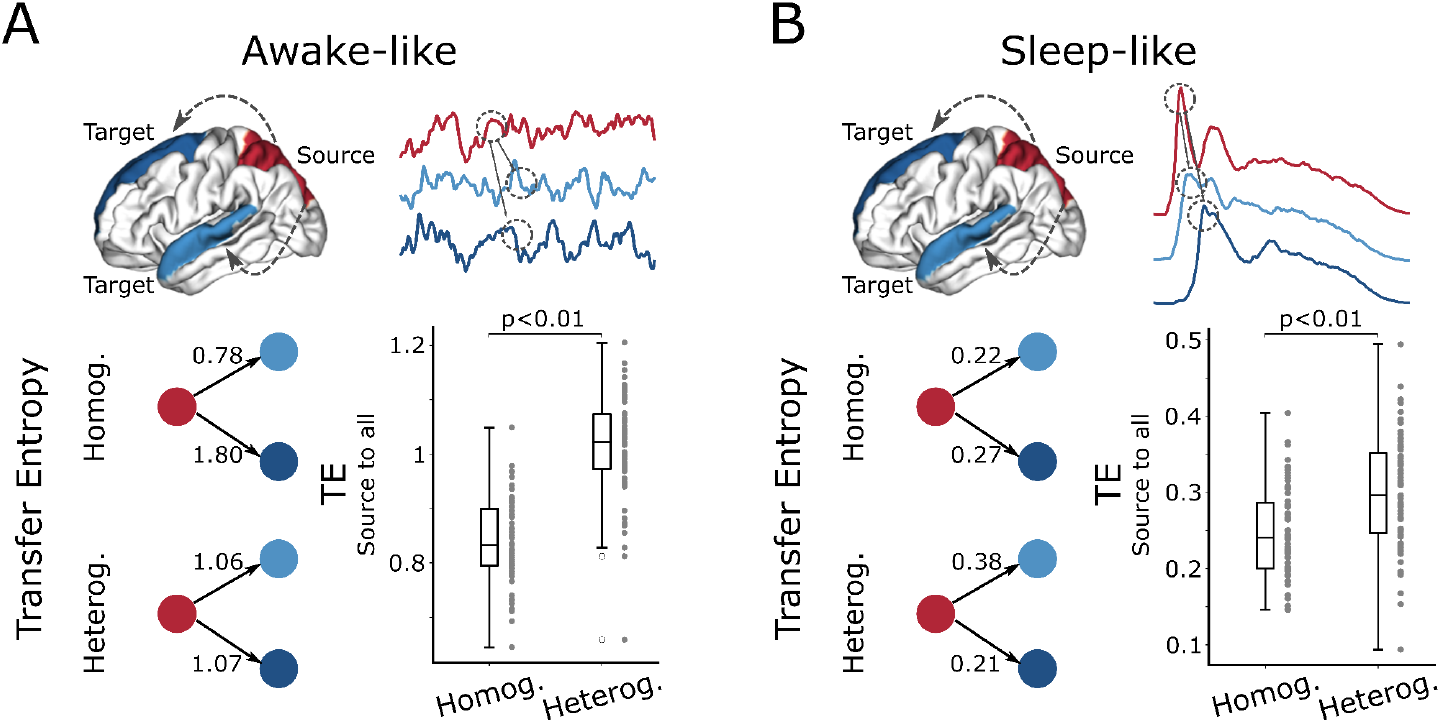
Information flow is enhanced in heterogeneous whole-brain networks. (A) Transfer entropy (TE) was computed from the stimulated area (source) to other brain areas during awake-like regime for heterogeneous (<b>=10pA) and homogeneous (b=10pA) networks. Bottom left illustrates TE between two example region pairs, highlighted in the brain illustration above. Bottom right shows the full TE profile from the source to all brain regions. (B) Same as in (A), but for the sleep-like regime, comparing heterogeneous (<b>=50pA) and homogeneous (<b>=50pA) networks. Statistical significance was evaluated using an independent two-sample t-test.

We found that, regardless of brain state, awake-like or sleep-like, TE was significantly higher in heterogeneous networks compared to their homogenous counterparts (Fig. 3). During the awake-like regime, when the network was more excitable and exhibited asynchronous dynamics, the mean TE (from source to all other areas) in heterogeneous networks was significantly higher than in homogeneous ones (1.02 ± 0.09 vs. 0.84 ± 0.08, respectively; Fig. 3A). Similarly, in the sleep-like regime, characterized by Up and Down dynamics, TE remained higher in the heterogeneous condition (0.30 ± 0.07 vs. 0.25 ± 0.06, respectively; Fig. 3B), suggesting that regional variability supports more efficient information flow even under globally synchronized states. Notice that TE values were also consistently higher in awake-like states compared to sleep-like states, in agreement with previous findings that Up and Down dynamics are associated with a breakdown in effective connectivity and a reduced capacity for information processing ^46–50^. Together, these results suggest that regional heterogeneity enhances information flow in large-scale brain networks.

### Emergence of localized sleep-like slow waves

We now return to the intermediate level of adaptation (Fig. 2, <b> = 25 pA). As previously described, this regime was characterized by the emergence of locally synchronized clusters in the FC, while the global network remained only weakly correlated (Fig. 2A-D and 4A). Here, we further examine the spatiotemporal features of this intermediate state and its underlying correlates (Fig. 4).

**Fig. 4.**
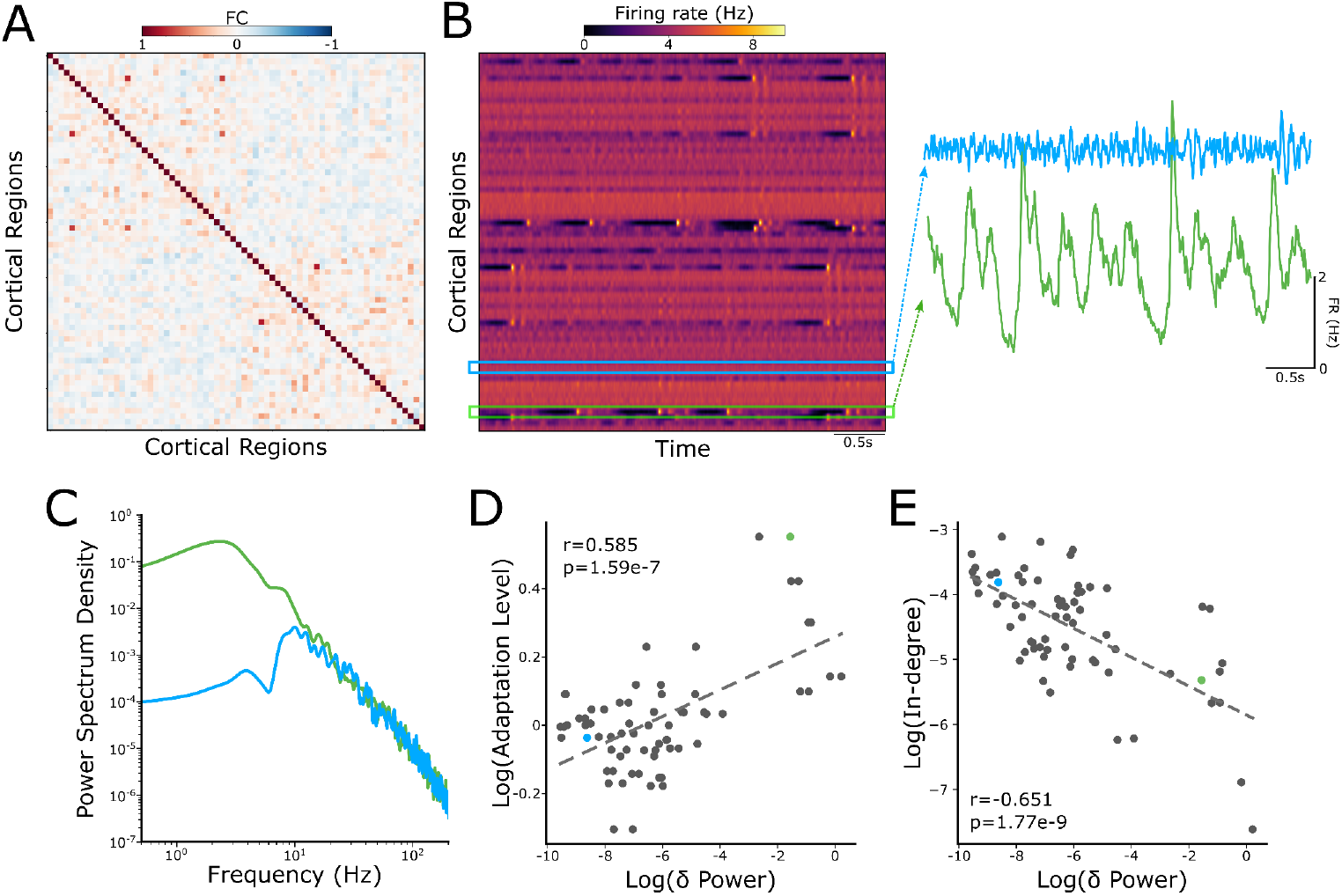
Emergence of localized sleep-like slow waves. (A) Functional connectivity (FC, Pearson correlation) matrix at an intermediate level of adaptation (<b>=25 pA). (B) Raster plot of network firing rates. Blue and green rectangles highlight the coexistence of two distinct dynamics: awake-like and sleep-like, respectively. Corresponding time series are shown on the right. (C) Power spectrum density (PSD) for the regions highlighted in (B). (D) Correlation between adaptation level and delta power (mean PSD power in the 0-4Hz band) across regions (r=0.585, p-value=1.59e-7). (E) Same as in (D) but showing the correlation between in-degree connectivity (average incoming input strength) and delta power (r=-0.651, p-value=1.77e-9). Pearson correlation coefficients and two-tailed p-values were used to assess statistical significance.

By plotting the firing rate of each brain region over time (Fig. 4B), we observed that few brain regions exhibited sleep-like slow waves, i.e., Up and Down dynamics, while others maintained a persistent level of activity, typically from awake-like states (Fig. 4B). These distinct temporal and spectral patterns, characterized by low-frequency, high-amplitude oscillations versus high-frequency, irregular fluctuations, coexisted across different regions within the same network (Fig. 4B-C), revealing an intricate dynamic state for intermediate levels of adaptation.

However, what determines whether a region expresses sleep-like or awake-like dynamics in this intermediate regime? To investigate this, we correlated the local adaptation levels with the delta power (mean power below 4 Hz; Fig. 4D). As expected, these two variables were highly positively correlated, indicating that regions with higher adaptation values were more prone to shift into the sleep-like mode. However, adaptation was not the only factor. The in-degree connectivity (average incoming input strength) also showed a strong negative correlation with delta power (Fig. 3E), suggesting that regions receiving fewer excitatory inputs were more prone to display sleep-like dynamics. Additionally, we observed a negative correlation, though statistically not significant, between adaptation levels and in-degree connectivity (r=-0.23; p=0.056). Together, these findings indicate that, in our model, localized sleep-like slow waves can emerge and coexist within an otherwise awake-like state, and that their spatial distribution is partially shaped by both regional adaptation levels and local connectivity properties. Moreover, our results are in line with recent empirical evidence that the level of regional cholinergic projections correlates inversely with the emergence of localized synchronous sleep-like activity ^17^.

## Discussion

Cortical dynamics are shaped by anatomical connectivity, as well as molecular and cellular composition, all of which vary across different brain regions ^6,7,40,51–54^. However, the extent to which this regional heterogeneity influences collective global brain dynamics has remained less explored, particularly across different brain states. In this study, we investigated the role of cholinergic regional heterogeneity using a whole-brain network model constrained by empirical human connectivity and transcriptomics data. Regional cholinergic heterogeneity was incorporated by modulating local node functional properties, thereby creating a more detailed virtual brain landscape (Fig. 1). Our results show that regional heterogeneity not only shapes spatiotemporal activity patterns by facilitating network synchronization but also enhances inter-regional communication. These findings underscore the significance of region-specific molecular profiles in shaping the dynamics of large-scale networks.

Brain states span a multidimensional space, from highly correlated (e.g., slow wave sleep) to weakly correlated (e.g., awake, attentive states) states ^2,3^. Transitions between these states are strongly influenced by neuromodulatory systems, which regulate neuronal excitability and network coordination ^5,55–58^. Among these, ACh plays a central role in shaping brain state dynamics by reducing spike-frequency adaptation, thereby increasing cortical excitability and modulating network synchrony ^5,13,59–63^. ACh levels also fluctuate according to the brain state, being elevated during wakefulness and REM sleep, and decreasing during slow wave sleep ^60,64,65^. Importantly, ACh release in the cortex, primarily stemming from the basal forebrain, is not spatially uniform, suggesting that neuromodulatory influence in neuronal dynamics may vary across brain regions ^15–17,66–68^.

Here, we aimed to investigate how this regional cholinergic heterogeneity shapes large-scale brain dynamics (Fig. 1). To this end, we employed the MF-AdEx model constrained by human structural connectivity derived from tractography data ^21,25,26^. This model has previously been shown to reproduce both awake-like and sleep-like brain state dynamics ^24,26^. Regional heterogeneity was introduced by modulating adaptation strength based on the gene expression levels of muscarinic acetylcholine receptors M1 (CHRM1) and M2 (CHRM2) ^6^. These receptor subtypes are two of the most abundant muscarinic receptors in the human cortex and are key targets of cholinergic modulation ^5,13,28,29,60^.

Our whole-brain model, incorporating cholinergic regional heterogeneity, reproduced key features of both awake- and sleep-like brain states (Fig. 2). During awake-like states, network dynamics were characterized by desynchronized patterns of activity and a suppression of low-frequency content. In contrast, sleep-like states were characterized by highly synchronous dynamics, dominated by propagating slow wave oscillations in the form of Up (active) and Down (silent) states (Fig. 2 and Supp). Additionally, the model also captured the brain state-dependent relationship between structural and functional connectivity (Fig. 2E). Specifically, during awake-like states, FC was only loosely constrained by the SC, whereas in sleep-like states, FC became more tightly aligned with SC. This strengthened FC–SC coupling has been observed in empirical human brain data and is thought to reflect a reduced repertoire of accessible dynamic states, a hallmark of unconscious states like deep sleep, compared to the broader dynamical flexibility present during wakefulness ^35–37,69–72^.

Beyond reproducing key features of brain states, what is the impact of regional heterogeneity in large-scale brain dynamics? Previous studies have shown that introducing spatially varying parameters such as local E:I balance, myelination gradients, or gene expression profiles significantly improves the fit of whole-brain models to empirical data ^10–12,73^. Heterogeneous models not only better replicate empirical brain activity but also support a richer dynamical repertoire, including more realistic regional timescales, ignition-like phenomena, and coherent spatiotemporal patterns ^11,12,74–76^. Thus, regional heterogeneity is essential for bridging anatomical specialization and global functional organization in computational models of brain activity.

Here, at the whole-brain level, we found that regional heterogeneity facilitates large-scale synchronization when compared to its homogeneous counterpart (Fig. 2D). Using the mean FC as a proxy of network state dynamics, we found that, as a function of adaptation levels, heterogeneous networks tend to exhibit higher synchrony. Additionally, the phase lag index, which captures non-zero-lag correlations, confirmed that heterogeneous networks tend to synchronize more easily (Fig. 2F). Although this effect has been previously reported in systems of heterogeneous oscillators ^77^ or spiking neurons ^45^, our results extend this concept to a biologically realistic whole-brain network model. These findings suggest that regional variability contributes to the emergence of coherent brain-wide activity, an essential feature of the brain implicated in a diversity of physiological and cognitive processes ^3,78–84^.

Another fundamental effect of regional heterogeneity observed in our model was its influence on inter-regional communication. Previous work shows that during awake-like states, information transfer is facilitated, while during sleep-like states, most probably due to off-periods, information processing is hindered ^46,47,49,50,72,85–90^. Our model was able to reproduce these empirical observations. Specifically, we showed that information transfer was greater during awake-like states compared to sleep-like states (Fig. 3). More importantly, we showed that regional heterogeneity significantly enhanced information flow when compared to homogeneous networks. This finding highlights the functional relevance of regional variability in supporting effective and flexible brain-wide communication. Moreover, these results align with previous research showing that heterogeneity improves the brain’s capacity for information transmission and processing ^11,43–45,91,92^.

Finally, we explored a special case: the coexistence of sleep-like slow waves within an otherwise awake brain. This complex phenomenon, where two distinct dynamic states coexist, has been previously reported under both physiological and pathological conditions. For example, in sleep-deprived awake rats, local slow waves have been observed within the awake brain and linked to impaired task performance ^93^. In humans, similar localized slow waves have also been observed during wakefulness, associated with attentional lapses under physiological conditions ^18^. Moreover, in both animals and humans, local slow waves have also been observed during physiological REM sleep ^17,20,94–96^. During pathological conditions, these localized slow waves have also been observed around brain lesions and epileptogenic zones ^19,97–101^. Thus, understanding the correlates of these local slow waves is crucial to understanding both physiological and pathological brain processes.

In our model, we reproduced this phenomenon, generating localized slow-wave activity within an overall awake-like dynamic state (Fig. 4). These localized slow waves were partially driven by regional heterogeneity in adaptation levels. Indeed, a recent study in animal models has also reported the occurrence of local slow waves during REM sleep, where their emergence has been associated with regional differences in cholinergic innervation ^17^. However, in our model, adaptation alone was not sufficient to fully explain the emergence of localized slow waves. The pattern of structural connectivity also played a key role, as it influenced the level of excitation each region received. This interplay is coherent with theoretical studies, where the level of adaptation and excitability are two key components of slow oscillations’ emergence and maintenance ^102,103^. Together, these results suggest that localized slow wave activity arises from a complex interplay between molecular and structural features of the brain. While we did not simulate pathological conditions, our framework could be extended to investigate how brain lesions and alterations in neurotransmitter levels might influence the generation of local slow waves after brain lesions ^19,104–106^, thus offering translational potential for developing personalized models of brain function and dysfunction.

In summary, our computational model highlights the significant impact of cholinergic heterogeneity on large-scale cortical dynamics. It introduces a novel framework for incorporating molecular-level variability into biophysically grounded whole-brain simulations. Although our approach relied on post-mortem transcriptomic data ^6^, future research could integrate other modalities such as receptor density maps obtained through positron emission tomography, especially since the relationship between these two data modalities is not yet clearly understood ^107^. Furthermore, exploring time-dependent neuromodulatory influences, such as how behavior-dependent fluctuations related to acetylcholine, or any other neurotransmitter associated with neuronal excitability, could be linked to the emergence of localized sleep-like activity, represents a promising and important direction for future work.

## Acknowledgments

Funded by INFRASLOW PID2023-152918OB-I00 financed by MICIU/AEI/10.13039/501100011033/FEDER, UE, by the European Union (ERC, NEMESIS, project number 101071900) and AGAUR co-funded by Departament de Recerca i Universitats de la Generalitat de Catalunya (AGAUR 2021-SGR-01165). Previously funded by the European Union’s Horizon 2020 Framework Programme for Research and Innovation under the Specific Grant Agreement No. 945539 (Human Brain Project SGA3). LDP acknowledges support from the Brazilian agency CNPq (Grant No. 140660/2022-4).

## Methods

### Mean-Field Model

The dynamics of each node were simulated according to a mean-field (MF) description of networks of Adaptive Exponential Integrate and Fire neurons (AdEx), as previously described ^21,24,26,108–110^. Briefly, this model captures the interaction among excitatory (E) and inhibitory (I) neuronal populations. Also, excitatory populations are equipped with firing rate adaptation (W) dynamics. The dynamics equations read:

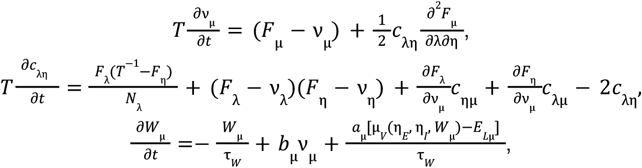

where *v*_µ_ is the mean firing rate of the population µ = {*E, I*}. *C*_λη_ is the covariance between population λ and η, *W*_µ_ is the mean adaptation, *b*_µ_ is the adaptation level (strength), and *a*_µ_ is the subthreshold adaptation, and *T* is the MF characteristic time constant. *F*_µ_ = *F*_µ_ (*v*_*E*_, *v*_*I*_, *W*_µ_) is the transfer function (TF), which characterizes the dependence of the output firing rate on the excitatory (*v*_*E*_) and inhibitory (*v*_*I*_) inputs. According to ^108^, it can be written as a function of its mean subthreshold membrane voltage µ _*V*_, its standard deviation *σ*_*V*_, and its time correlation time decay τ _*V*_, as:

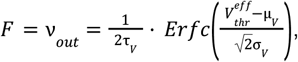

where (µ _*V*_, *σ* _*V*_, τ _*V*_) are obtained by solving a set of equations, as described in ^21,108^.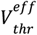 is a phenomenological spike threshold voltage taken as a second-order polynomial:

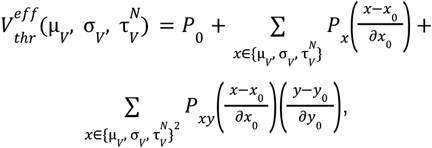

with 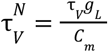, where *g*_*L*_ is the leakage conductance and *C*_*m*_ the membrane capacitance. The constant values are defined as in ^24^:µ_*V*_ =− 60 *mV, σ*_*V*_ = 0. 004 *mV*, 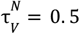, dµ_*V*_ = 0. 001 *mV*, d*σ*_*V*_ = 0. 006 *mV*, and 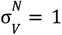. Accordingly, the fitted polynomials *P*^*E,I*^, for *E* and *I* types of neurons are: *P*_0_ =− 0. 0498, − 0. 0514, *P*_µ*V*_ = 0. 00506, 0. 004, *P*_*σV*_ =− 0. 025, − 0. 0083, *P*_τ*V*_ = 0. 0014, 0. 0002,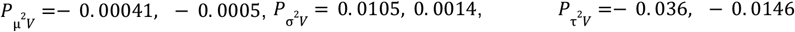 *P*_µ*VσV*_ = 0. 0074, 0. 0045, *P*_µ*V*τ*V*_ = 0. 0012, 0. 0028, *P*_*σV*τ*V*_ =− 0. 0407, − 0. 0153, for (*E, I*), respectively. µ_*V*_ is a function that represents the average membrane potential of a given population, described by:

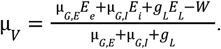

µ _*G,E*_ = *v*_*E*_ *K*_*E*_ *u*_*E*_ *Q*_*E*_, and the same applies to the *I* population. *Q* is the conductance weight, *u* is the synaptic time decay, and *K* = *Np* is a constant defined by the number of neurons (*N* = 10^4^) and the probability of connection (*p* = 0. 05). All parameters were obtained from and are set to: *T* = 20ms, *E* _*L*,{*E,I*}_ = (− 64, − 65)*mV, N*_{*E,I*}_ = (8000, 2000), *p* = 5%, *b*_*I*_= 0*pA, a*_{*E,I*}_ = (0, 0)*pA*, τ_*W*_ = 500*ms, g*_*L*_ = 10*nS, C*_*m*_ = 200*pF, Q*_{*E,I*}_ = (1. 5, 5)*nS, v*^ext^ = 0. 315*Hz, K* _{*E*.*I*}_ = (400, 0), *E* _*e,i*_ = (0, − 80). *b*_*E*_ sets the adaptation level of the excitatory populations and was used as a proxy of spatial cholinergic heterogeneity; see the *Regional Heterogeneity* section below.

The previous equations describe the population dynamics of a single cortical region composed of excitatory and inhibitory populations. To describe large-scale networks of interconnected brain regions, each represented by the MF model, we can extend the TF to incorporate both inter-regional interaction and external noise. Thus, the TF can be rewritten as simply as:

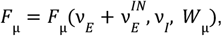

with,

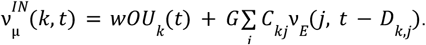

*K* represents the node index, *G* = 0. 3 is the global coupling factor that scales all the connection weights, *C*_*kj*_ is the connectivity matrix strength between *j* and *k, D*_*k,j*_ is the matrix delay of axonal propagation, computed as ∥*j* − *k*∥/*v* _*c*_, where ∥*j* − *k*∥ is the distance between nodes *j* and *k* and *v*_*c*_ = 5*mm*/*ms* is the propagation speed, *w=1e-4pA* is the noise scaling factor, and *OU* _*k*_(*t*) is the noise defined by an Ornstein-Uhlenbeck process with τ_*OU*_ = 5*ms*.

#### Structural Connectivity

The structure connectivity (SC) was derived from human tractography data from the Berlin empirical data processing pipeline ^25^. A cortical parcellation of 68 brain regions was used, according to the Desikan-Killiany atlas. Connection strengths (weights) and transmission delays were based on estimates of fiber density and tract length, respectively, obtained from diffusion tensor imaging (DTI) data ^27^. The preprocessed data was obtained from ^25^, and was the same used in a previous study ^26^.

#### Regional Heterogeneity

Regional heterogeneity was implemented by incorporating spatial variation in cortical cholinergic receptor density to modulate the level of neuronal adaptation in the MF-AdEx model. Gene expression data were obtained from the Allen Human Brain Atlas (AHBA) and were processed according to the ABAGEN toolbox using robust sigmoid normalization and RNA-sequencing-based probe selection, following established protocols for minimizing inter-donor variability and preserving spatial fidelity ^6,11,30,31^. Departing from our knowledge on the role of cholinergic modulation in cortical circuits ^13^, we focused on muscarinic receptor genes. Specifically, muscarinic acetylcholine receptor subtypes M1 (CHRM1) and M2 (CHRM2), the two most abundant muscarinic receptors in the human cortex ^6,28,29^. Because AHBA data contain more samples from the left hemisphere, we extracted expression values from the left cortex and mirrored them to the right hemisphere, consistent with previous modeling approaches ^11^.

In our implementation, we used transcriptomic data from the cholinergic muscarinic receptors CHRM1 and CHRM2. As their density were highly correlated across regions (see Fig. 1 bottom panel), we worked with the normalized arithmetic mean, which we refer to as simple CHRM density. To ensure equivalence with a homogeneous network, we centered this distribution around its median, thus centering it around one (1.03±0.20, mean and standard deviation given). The resulting CHRM density vector, containing 68 values (one per cortical region), was then used to modulate the adaptation parameter *b*_*E*_ as:

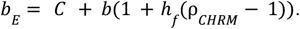

*C* = 10*pA* is the baseline adaptation level, *b* is a free variable that represents the adaptation. ρ_*CHRM*_ represents the CHRM density, and *h*_*f*_ is the heterogeneity factor, i.e., if *h*_*f*_ = 1 the network is fully heterogeneous, while *h*_*f*_ = 0 represents a fully homogeneous network.

#### Simulations

The model was run using The Virtual Brain (TVB) platform, where the MF-AdEx model is already implemented ^111,112^. Numerical integration was performed using the stochastic Heun method with a time step of 1 ms. For spontaneous activity simulations, each run lasted 22 s, with the first 2 s discarded to eliminate transients. For stimulation protocols, a brief excitatory input was applied to the firing rate of the excitatory population in a single region, the superior parietal lobule. The stimulus had a duration of 30 ms and an amplitude of 3 Hz. Each stimulation trial (50 trials) consisted of a new simulation lasting 6.03 s, with the first 2 s again discarded. The stimulus was applied at 4 s, allowing for a 2 s pre-stimulus and 2 s post-stimulus window for analysis.

#### Data Analysis

Functional connectivity (FC) and structure-function coupling (SC-FC) were estimated using the Pearson correlation. Power spectrum density was computed using Welch’s method. Phase Lag Index (PLI) was computed as described in ^39^: *PLI* = |< *sign*[Δϕ(*t*)] >|. Where Δϕ denotes the instantaneous phase difference between two signals. The instantaneous phase was estimated by the Hilbert transform. PLI measures the phase synchronization among signals while minimizing the influence of zero-lag correlations. PLI values range from 0 and 1, where 0 indicates no consistent phase lag, and 1 reflects perfect phase locking with a consistent non-zero phase difference ^113^. Transfer entropy (TE) between two signals *X*(*t*) and *Y*(*t*), *TE* _*X*→*Y*_ was defined as: *TE*_*X*→*Y*_ = *H*(*Y*_*t*_ |*Y*_*t*−1:*t*−τ_) − *H*(*Y*_*t*_ |*Y*_*t*−1:*t*−τ_, *X*_*t*−1:*t*−τ_), where *H*() denotes the entropy and τ _*X*→*Y*_ = 5*ms* is the time delay. In short, *TE* measures the reduction in uncertainty about the future of *Y*(*t*) given its own past and the history of a second variable *X*(*t*) ^114,115^. We used the TE implementation available at: https://github.com/notsebastiano/transfer_entropy. TE was computed from the source area to all other areas in a window of 400 ms after the stimulus (see above).

## Code and Data Availability

The MF-AdEx code used in this study is freely available through The Virtual Brain (TVB) platform (https://www.thevirtualbrain.org/). The transcriptomic data utilized are publicly accessible via the Allen Institute for Brain Science at https://human.brain-map.org/.

---

**Supplementary Fig.**
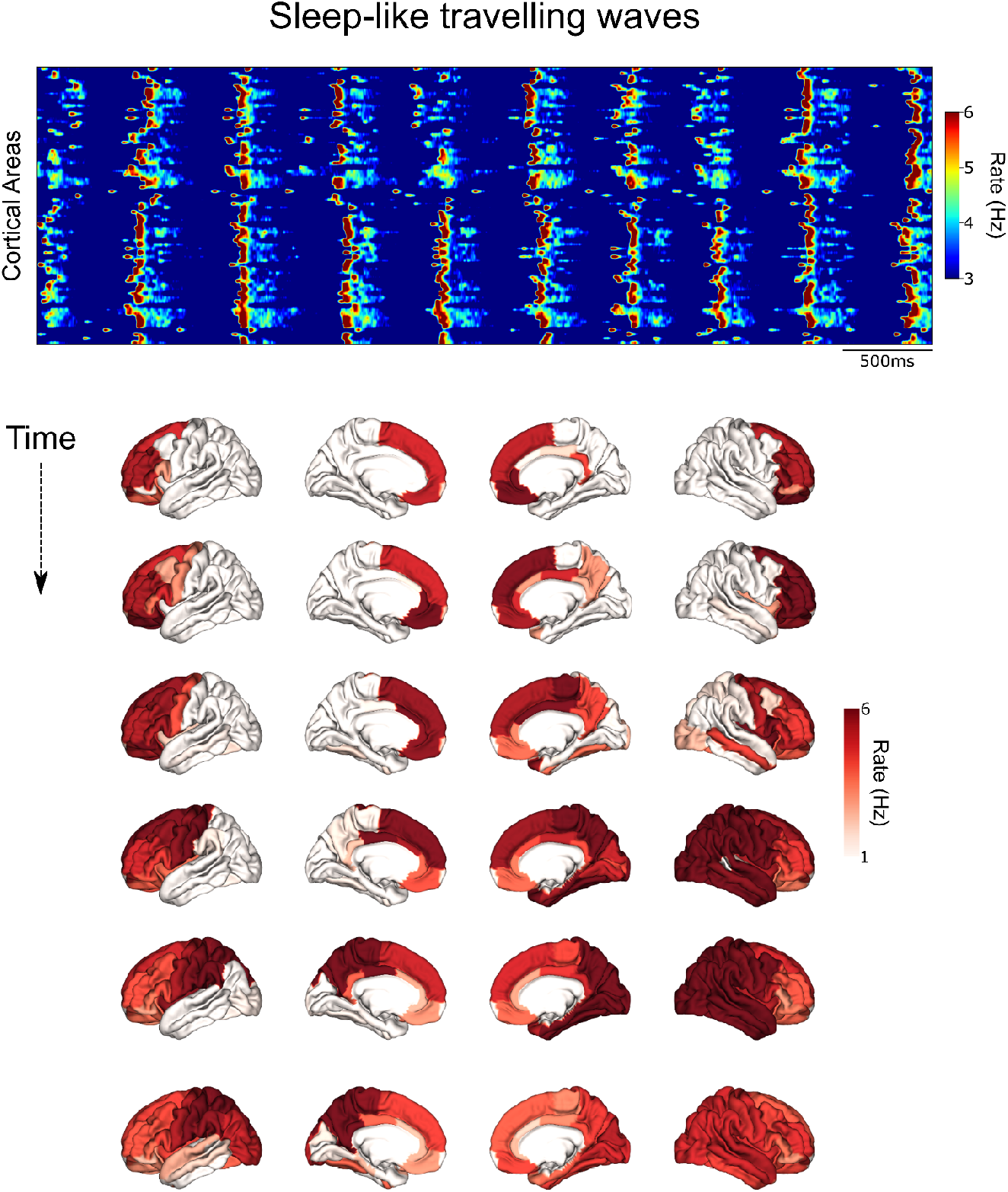
Cortical travelling slow waves. Top, Raster plot of cortical areas illustrating Up and Down dynamics and wave propagation. Bottom, Illustration of a travelling wave during one Up state. Each row is separated by 20 ms.

